# Microscopy-guided subcellular proteomic discovery by high-speed ultra-content photo-biotinylation

**DOI:** 10.1101/2023.12.27.573388

**Authors:** Yi-De Chen, Chih-Wei Chang, Chantal Hoi Yin Cheung, Hsiao-Jen Chang, Yong-Da Sie, Chia-Wen Chung, Chun-Kai Huang, Chien-Chang Huang, Weng Man Chong, You-Pi Liu, Yu-Chih Lin, Hsiang-Ju James Kai, Pei-Jie Wang, Jung-Chi Liao

## Abstract

Microscopy-guided proteomics at an organelle-dimension resolution is desired for revealing unknown protein constituents at specific disease- or functional-associated regions at the molecular-molecular interactions level. Here, we achieve protein spatial purification by introducing a firmware-integrated microscopy platform that triggers *in situ* subcellular photo-biotinylation of proteins at user-defined regions of interest (ROIs) one field of view (FOV) at a time for thousands of FOVs fully automatically. An illumination pattern at the analogous ROIs of each FOV is calculated on the fly by either machine learning or traditional image processing. Photoactivatable amino acid crosslinkers are activated by a two-photon focal light one spot at a time at a sub-millisecond illumination duration per spot. Imaging, pattern generation, targeted illumination, and FOV movement are coordinated and cycled with high-speed mechatronic control to complete illumination on millions of ROI spots within hours. Once enough proteins are biotinylated in a cell or tissue sample, the sample is scraped and lysed, and avidin pulldown is used to enrich proteins to achieve spatial protein scooping at a 240-nm precision. Subsequent LC-MS/MS is implemented to reveal the subcellular proteome in high sensitivity, specificity, and resolution. Using this technology termed optoproteomics, we have revealed novel stress granule-localized and amyloid β-localized proteins validated by immunostaining. Together, spatial purification by ultra-content, high-speed microscopy-targeted photo-biotinylation enables unprecedented subcellular spatial proteomics discovery in any microscopically recognizable regions.

## Introduction

Pathology, neuroscience, oncology, cell biology, and other life science fields use microscopy to recognize, localize, or map features of interest, where it is usually challenging to physically isolate proteins *in situ* and identify them at a specific subcellular region of interest (ROI) that represents a disease signature or corresponds to important molecular-molecular interactions. For example, it would be desired to physically pull-down proteins from thousands or millions of microscopy-recognized immune synapses, TDP-43 aggregates, focal adhesions, special-shaped dendritic spines, tips of primary cilia, insulin granules, or mitochondria-lipid droplet contact sites automatically and use mass spectrometry to determine the exact protein constituents at these sites. The lack of this unbiased organelle-scale spatial proteomic discovery capability hinders identification of novel disease-associated proteins, especially when they are functionally localized or relatively low abundant.

Spatial proteomics allows protein mapping of a biological sample to reveal geographic organization for underlying protein-protein interactions (*1–3*). Targeted spatial proteomics mostly uses known antibody arrays aiming to localize known proteins (*4, 5*), whereas *de novo* spatial proteomics requires spatial protein identification without prior knowledge of what proteins to look for. Unlike transcriptomics where PCR is capable of amplifying signals so that *de novo* transcriptomics like RNAseq is possible, no PCR-equivalent technology is yet available for proteomics.

Two major techniques are feasible for *de novo* spatial proteomics: large-scale microscopy mapping and mass spectrometry (MS). The Protein Atlas Project mapped thousands of protein species using large-scale immunostaining, effectively creating a *de novo* spatial proteomic database (*6*). The limitation of this approach is its applications to specific biological problems, where a multi-month extensive staining process must be implemented for each project if the proteome variation due to a stimulation, a knockdown/knockout, or a mutation is of interest. Immunoprecipitation (IP) and MS together is a widely used biochemical approach to identify a proteome associated with a bait protein. Recent proximity labeling approaches provide further improvement in spatial precision close to a bait protein (*7–12*). These are broadly applicable for diverse biological problems when the biochemical isolation is specific. For some applications when non-specific interactions of the pulldown process for IP and proximity labeling introduce undesired proteins in the proteome, the specificity of the methods will be impaired. In the case when an ROI can only be specified by morphology characteristics without an apparent bait, proximity labeling approaches will not work. For animal and human samples, expression of an exogenous enzyme for proximity labeling requires significant transfection or knockin efforts (*13–18*). Laser capture microdissection (LCM) including the recent AI-guided deep visual proteomics (DVP) enables protein isolation at specific ROIs and subsequent *de novo* spatial proteome identification (*19–21*). DVP provides a powerful improvement in throughput for spatial proteomic discovery, whereas the beam diameter of the cutting laser remains in a cutting speed-dependent micron size, requiring extra efforts for subcellular dissection precision. Its non-discriminative vertical cutting collects non-specific proteins in different axial planes and may affect specificlity. Recent development of spatially targeted optical microproteomics (STOMP) and its derivative approaches such as autoSTOMP offer another *de novo* spatial proteomics technique to identify the proteome at specific ROIs under a microscope (*22–24*). However, without a hardware change to speed up the process, it needs a long experimental duration to reach the fundamental scaleup requirement for MS sensitivity and specificity due to the ROI illumination speed, making it feasible for validating known high abundant proteins but difficult to fulfill the major discovery goal, where sensitivity is required to unveil relatively low abundant proteins that are unknown for a specific biological problem.

Here, we developed a novel technology termed optoproteomics, a proteomic discovery approach enabled by fully automated and synchronized microscopy system that performs high-speed ultra-content *in situ* protein photo-biotinylation. Mechatronic integration of multiple processing steps at a synchronized sub-millisecond time scale was implemented to assure manageable duration of hours for the entire process of targeted illumination on millions of diffraction-limited spots. Subsequent biochemical pulldown and nano-LC-MS/MS were implemented for protein enrichment and proteome identification at the specified subcellular ROIs. This optoproteomics method overcomes the protein amplification issue by targeted protein accumulation and achieves *de novo* subcellular proteomic discovery in high sensitivity, specificity, and resolution.

## Results

### Speed optimization is essential for vast spatial protein purification and accumulation

To achieve high spatial resolution and high specificity of microscopy-guided proteomics, two-photon-induced protein biotinylation at selected ROIs, one spot at a time through mechatronic position control, as well as subsequent biochemical pulldown and nano-LC-MS/MS were implemented, with detailed photochemistry described below. The two-photon illumination was used because it achieved better chemical labeling precision in the axial direction and negligible scattering-induced photochemical reaction in 3D. For MS-based proteomics, roughly ∼10^8^ molecules of the same species of proteins are needed for the sensitivity of routinely used MS platforms in a manageable dynamic range (*25, 26*). To achieve high sensitivity of *de novo* subcellular proteomic discovery, one should collect proteins from myriad ROIs of a large number of fields of view (FOVs) with analogous protein constituents. For a relatively low copy number protein such as one with 1,000 copies per cell, and assuming the photo-biotinylation efficiency is 10%, one needs to illuminate 10^6^ cells for MS sensitivity. Assuming a FOV of a 40X and 0.95 numerical aperture (NA) objective that accommodates 50 cells, one should illuminate 2 × 10^4^ FOVs. In terms of the total number of illumination spots, approximately 0.1 µm^2^ is covered per diffraction-limited spot for a 40X NA 0.95 objective. The areas of ROIs largely vary depending on problems, from 2 µm^2^ per cell (i.e., 20 illumination spots) for primary cilia to ∼20 µm^2^ per cell (i.e., 200 illumination spots) for nucleoli. That is, a total of 2 × 10^8^ illumination spots is needed for nucleolar protein labeling. If one spot takes 0.1 sec for illumination, shutter on/off, and spot-to-spot movement, and one FOV takes 5 sec for imaging turret movement, focusing, pattern generation calculation, and stage FOV movement, one will need a total of 2.01 × 10^7^ sec, or ∼230 days to collect enough protein samples, too long to accomplish in a regular basis.

Here an ultra-content high-speed real-time targeted microscopy-based illumination system was developed to significantly expedite the illumination speed. Photolabeling on functionally analogous ROI locations was performed, under the assumption that major protein constituents in specific functional sites are largely similar. These functional sites are the sites whose morphological features or image contrasts are recognizable under a microscope. Precision photochemical reactions assured targeted protein labeling with a low background so that high-specificity microscopy-guided proteomics became feasible. There were four sequential steps repeating tens of thousands of times needed for the entire process: 1. microscopy imaging at an FOV; 2. ROI pattern generation of a FOV image on the fly by traditional image processing or deep learning; 3. directed illumination toward the ROIs for photochemical labeling; and 4. FOV change to the next one (Fig. 1a and Video 1-4). This repetitive process effectively performed protein spatial purification, serving as the core to achieve protein accumulation that overcame the fundamental problem of protein amplification. Existing technology or system was not optimized to perform this process repeatedly for so many times within a few hours. Without such a speed, one will only be able to identify high-abundant proteins, which are mostly known and lack novelty.

**Fig. 1.**
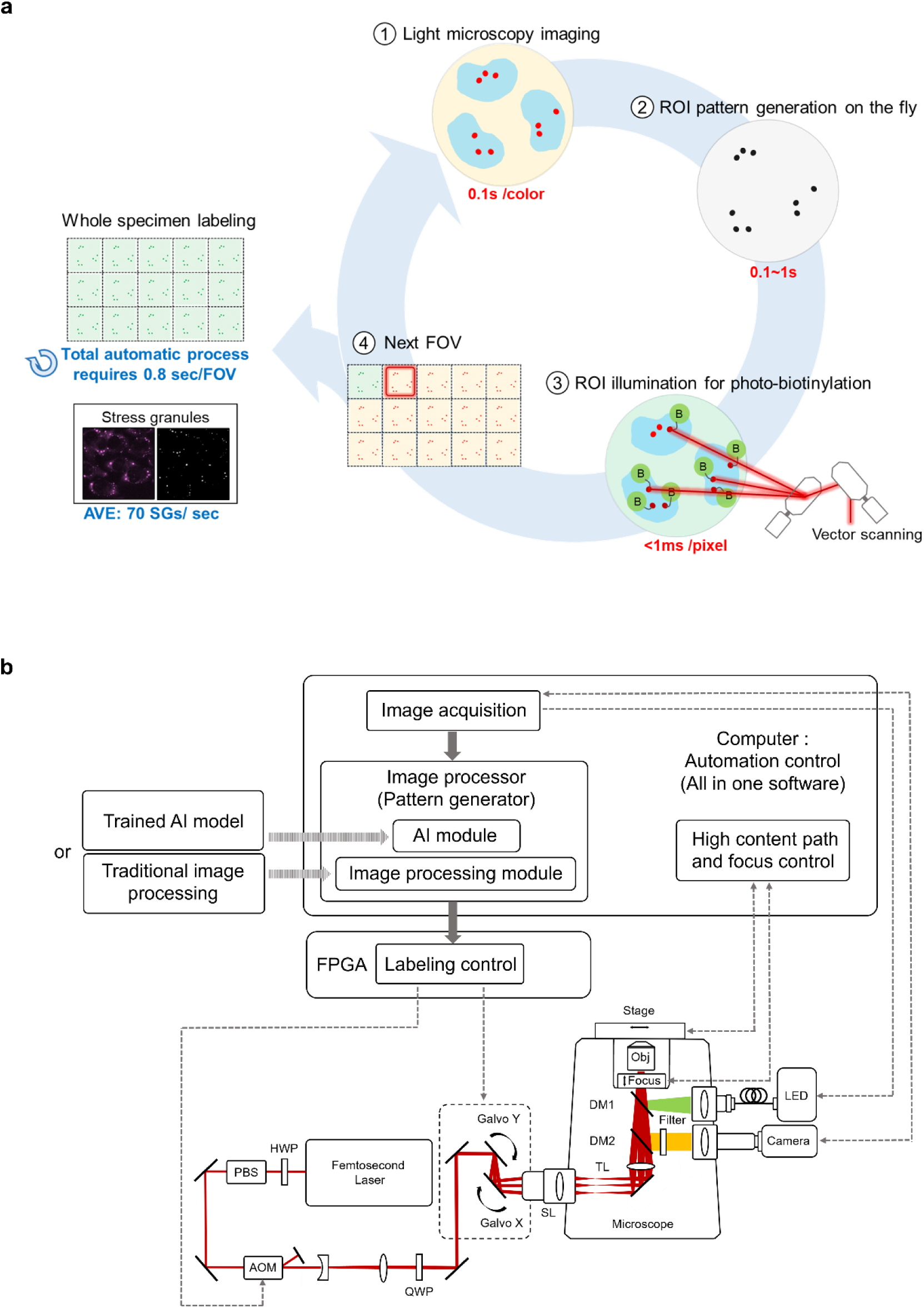
The process and design of the microscopy-guided protein purification system. **a,** Schematic overview of the workflow for ultra-content targeted photo-biotinylation. The process includes: (1) light microscopy imaging of a field of view (FOV); (2) pattern generation of the regions of interest (ROIs) on the fly; (3) precision illumination on the selected ROIs for protein photo-biotinylation; (4) stage movement to a next FOV; and (5) repeat of the steps 1-4 for each FOV until all FOVs of interest are processed. **b,** The optical setup and its control diagram of the spatial purification microscopy system. Two light sources are included in the system: an LED light source for imaging and a femtosecond laser light source for two-photon photolabeling. The femtosecond laser beam was directed by a pair of galvanometers to illuminate the sample at selected points of interest (see Fig. S1 for details). FPGA is used to synchronize the control of the fast galvanometer movement and acousto-optic modulator (AOM) shuttering. Images acquired by the camera are processed in real time by computer software with either traditional image processing or deep learning to generate the patterns of ROIs. The coordinates of the patterns are used to configure the FPGA for labeling control. DM, dichroic mirror; TL, tube lens; SL, scan lens; QWP, quarter-wave plate; PBS, polarizing beam splitter; HWP, half-wave plate.

The optical system in this study contained a motorized inverted light microscope with a drift-free focusing setup, an LED light source for multicolor fluorescence imaging (e.g., 488 nm, 568 nm, 647 nm), an sCMOS camera for imaging, a 780-nm femtosecond light source for two-photon illumination that triggers a photochemical reaction, an acousto-optic modulator (AOM) as a femtosecond light shutter, and a pair of galvanometer scanning mirrors (galvos) to direct the femtosecond light illumination (Fig. 1b and Fig. S1). Electronically switched LED and AOM minimized color switching speed at the time scale of <20 µs for LED triggering and 25 ns time scale for AOM rise/fall. To avoid any slowdown due to mechanical movement, multiband dichroic mirrors were used to allow multicolor imaging and femtosecond light illumination without movement of mechanical elements such as a turret or a shutter. FPGA with a 1-µs conversion time was used to control AOM and galvos to assume synchronization for scanning, where 100-500 µs per illumination spot was needed for the duration of a photochemical reaction illumination and the dwell time optionally required for galvo movement stabilization when labeling high-precision objects. The only mechanical movements required in the process were the fast galvo scanning and the relatively slower motorized focal adjustment and the stage movement toward the next FOV.

A software-firmware integrated program was required to control imaging, pattern generation calculation, photochemical illumination, and field change in a tight coordination. 100-ms exposure time was used for widefield imaging of each color when possible. The images were analyzed on the fly to segment the ROIs based on the user’s interest using either traditional image processing or deep learning in the software (Fig. 1b). When using traditional image processing, an image processing set of steps with a combination of methods such as thresholding, filtering, or other approaches was applied identically to all FOVs (Fig. 2a and Fig. S2). Some segmentation tasks were more complex than others, and pre-processing or post-processing might be added to normalize the image quality of different FOVs. Example segmentation results are shown in Fig. 2b. This image processing step took 0.1 to 1 sec depending on the processing complexity and image quality. Coordinates of all the grid points at the ROIs were then obtained. An optimized planned path was calculated and used to direct the galvos to scan through these grid points. The galvos and the AOM were synchronized at the ∼100 µs scale to allow on and off labeling at the right location with an exact illumination dwell time per spot to assure uniform photochemical reaction duration (Fig. 2c). For multiple ROIs, the scanning path went through one ROI at a time from an initiation point of the peripheral spiraling clockwise toward the center before moving to the initiation point of an adjacent ROI to minimize the traveling time as well as to reduce motion jerk for galvo damage protection (Fig. 2c). For a 100-500-µs illumination per spot, 10^7^-10^8^ illumination spots, 1-2 sec per FOV, and ∼10^4^ FOVs for a 2 cm × 4 cm sample well of ∼5 × 10^5^ cells under a 40X NA 0.95 objective, one needs 1.1-7 × 10^4^ sec, or 3-19.5 hr to complete one photolabeling round. Depending on the problems and the needed level of retrieving low abundant proteins, one to ten sample wells were needed to collect enough proteins for MS analysis. Speed optimization of every step, full automated synchronization, and mechatronic control made the *de novo* spatial proteomics feasible to finish within a reasonable time.

**Fig. 2.**
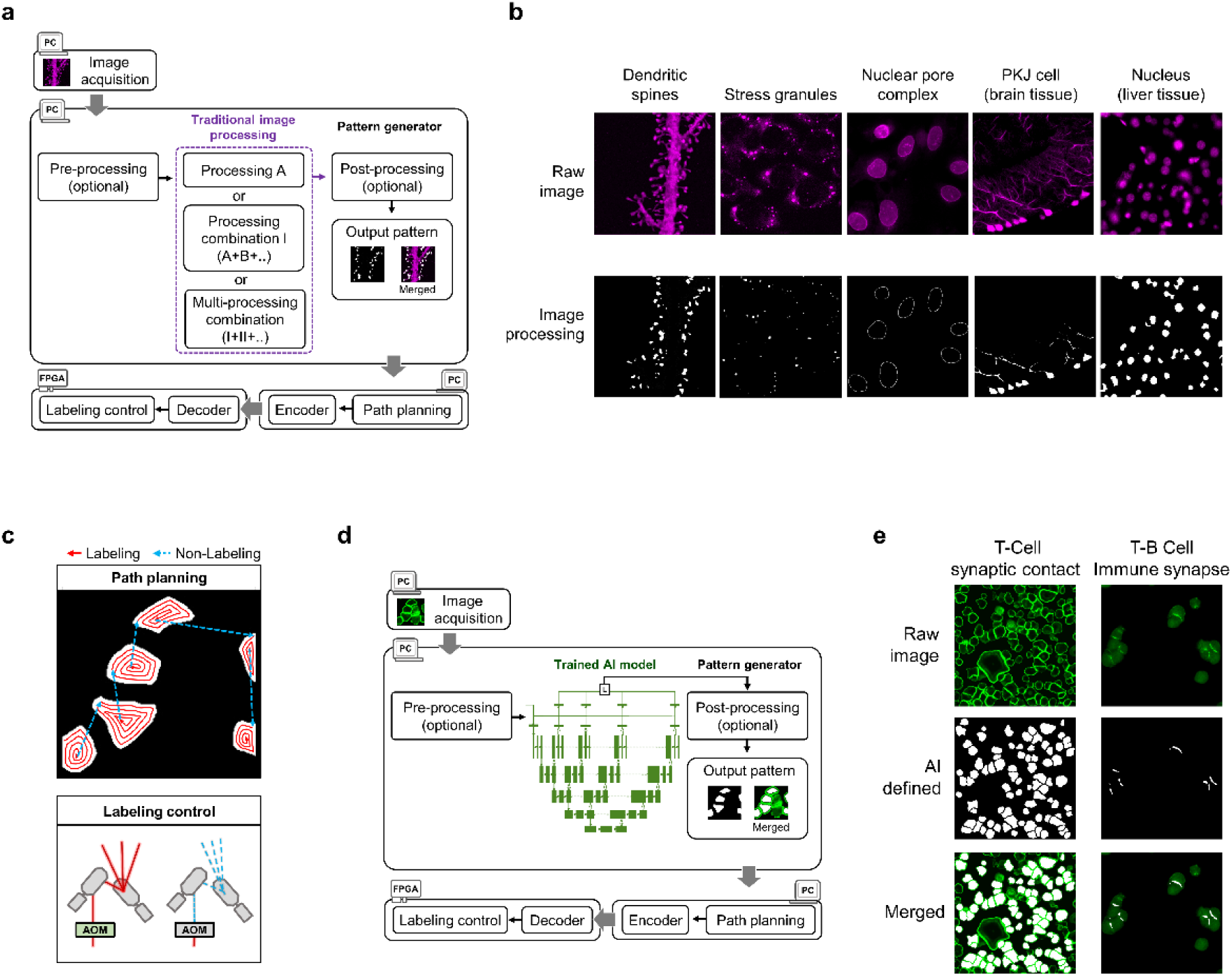
Automatic pattern generation for targeted photo-biotinylation by either traditional image processing or deep learning. **a,** Traditional image processing and associated steps to determine the regions of interest (ROIs) on the acquired images. Path planning is performed for all points of the generated pattern, and the array of the point coordinates is used for labeling control using FPGA. **b,** Patterns generated for various subcellular structures by traditional image processing. **c,** Paths of clockwise spiraling from the peripheral region are used to minimize the traveling time and reduce motion jerk. AOM is used to control illumination on and off. **d,** Implementation of deep learning for image segmentation of the ROIs. A U-Net model is used for training. e, Example patterns generated by deep learning.

### Deep learning enables better pattern generation

When images were more complex, or image quality was worse, deep learning-based image segmentation showed better accuracy in ROI pattern generation. A hundred of annotated images were used to train a deep learning semantic segmentation convolution neural network (Fig. 2d and Fig. S3), which was modified from U-Net with pre-trained weights from ImageNet to facilitate convergence (*27, 28*). The RMSprop optimizer was utilized to converge the dice coefficient loss function (*29, 30*). A heavier weight was assigned to false positives of the loss function at different resolutions of the model output to minimize false positive photolabeling. After multiple training epochs, we assessed the stability and accuracy of the model using a modified metric, which was also calculated at different resolutions of the model output.

The trained network was applied identically to each FOV, so that the image of each FOV was segmented on the fly based on this trained network. The grid points of the ROIs of an FOV were obtained and then path planning as well as labeling control were implemented (Fig. 2d). Two examples of segmentation by deep learning are shown in Fig. 2e.

### Photoactivatable amino acid crosslinking enables spatial photo-induced biotinylation

To achieve photo-induced labeling of proteins at a microscopic illumination spot, the sample was added with molecules containing three elements: a photocatalysis, a photoactivatable amino acid linker to covalently bind to a protein, and a probing tag for protein pulldown. One combination was to use biotin-phenol (BP) and a photosensitizer such as Rose Bengal, riboflavin, or [Ru(bpy)]_32+_, where the photoactivated photosensitizer triggered a photoredox reaction to oxidize the phenol functional group of BP molecules in the media and generated phenoxyl radicals, which quickly formed phenol-tyrosine coupling with neighboring proteins, resulting in *in situ* biotinylation of proteins near the illumination spot. Another way was to use biotin-benzophenone (BBZP). Benzophenone at the illumination spot was excited to become 1,2-diradical, which reacted with the C-H bond of α-carbon, forming a covalent bond with the corresponding amino acid and resulting in targeted biotinylation of proteins near the illumination spot.

There were two major challenges of photochemistry: reaction efficiency and spatial labeling specificity. In addition to fast galvo movement, the photochemical reaction also needed to be completed in as fast as 0.1-0.5 ms per spot to achieve ultra-content labeling. The spatial labeling specificity was essential because the ROIs might only occupy 1/10,000 or even smaller volume of the entire biological sample space. A tiny fraction of non-specific binding at the non-illuminated regions would result in a sizable collection of unwanted proteins, potentially overshadowing the accumulated copies of desired proteins at the ROIs.

We thus tested different photosensitizers, redox molecules, their concentrations, and illumination intensities to satisfy the speed and specificity needs of photochemistry. Among [Ru(bpy)_3_]^2+^, riboflavin, Rose Bengal, and Irgacure together with desthiobiotin-phenol (DBP, desthiobiotin instead of biotin was used to make the later elusion process easier), [Ru(bpy)_3_]^2+^ gave the best reaction efficiency when illuminating the entire FOV of U-2OS cells with a 780-nm femtosecond laser, illustrated by Dy488-NeutrAvidin imaging (Fig. S4a). The wavelength of 780 nm was chosen because of its excitation efficiency (Fig. S4b). 100-mW laser power at the back focal plane of a 40x 0.95 NA objective was enough for labeling. Addition of methyl viologen further improved the labeling efficiency (Fig. S4a). When using BBZP, labeling efficiency was similar to that of [Ru(bpy)_3_]^2+^ and DBP, but the non-specific labeling background was lower (Fig. S5), demonstrating that BBZP was a better choice to optimize both reaction efficiency and spatial labeling specificity.

### Subcellular spatial biotinylation is applicable to cell and tissue samples

To facilitate subcellular protein isolation, the labeling resolution should be at a submicron level. Illumination of single lines was performed on U-2OS cells and super-resolution structured illumination microscopy was used to measure the biotinylation labeling width (Fig. 3a). We were able to reach 240-nm labeling resolution when using a 40x 0.95 NA objective (Fig. 3b). To demonstrate its broad usage on cell samples, we photolabeled proteins at five subcellular locales including nuclei, nucleoli, nuclear pore complexes, stress granules, and Golgi apparatuses. Image segmentation was performed based on the characteristics of each structure using traditional image processing. The segmented region was illuminated to induce targeted biotinylation. The *in situ* biotinylated regions matched well with the corresponding subcellular structures in lateral (xy) and axial (z) directions, suggesting a high spatial labeling specificity (Fig. 3c). We also applied deep learning to segment the region within F-actin rings of Jurkat T cells in a spreading assay on an anti-CD3 coated glass surface. Segmented illumination at a focal level within the glass allowed photolabeling at a thin layer of the contact surface (Fig. 3d), allowing specific biotinylation of proteins at the supramolecular activating complexes (SMACs) of the immune synapses. Deep learning-based segmentation and targeted illumination were also applied to conjugates of Jurkat-Raji co-culture cells with precise biotinylation of proteins at thin immune synapses (Fig. 3e), illustrating the capability of photolabeling upon multi-color imaging. We further examined the applicability of photolabeling on tissue samples. Calbindin-D28K staining of a FFPE mouse brain tissue section revealed a high abundance of Purkinje cells in cerebellum with clear axon and dendrite architecture and a distinctive neuron cell body layer (Fig. 3f). Deep learning-enabled segmentation and following targeted illumination resulted in biotinylation at cell bodies (Fig. 3f), demonstrating photolabeling specificity for FFPE tissue samples.

**Fig. 3.**
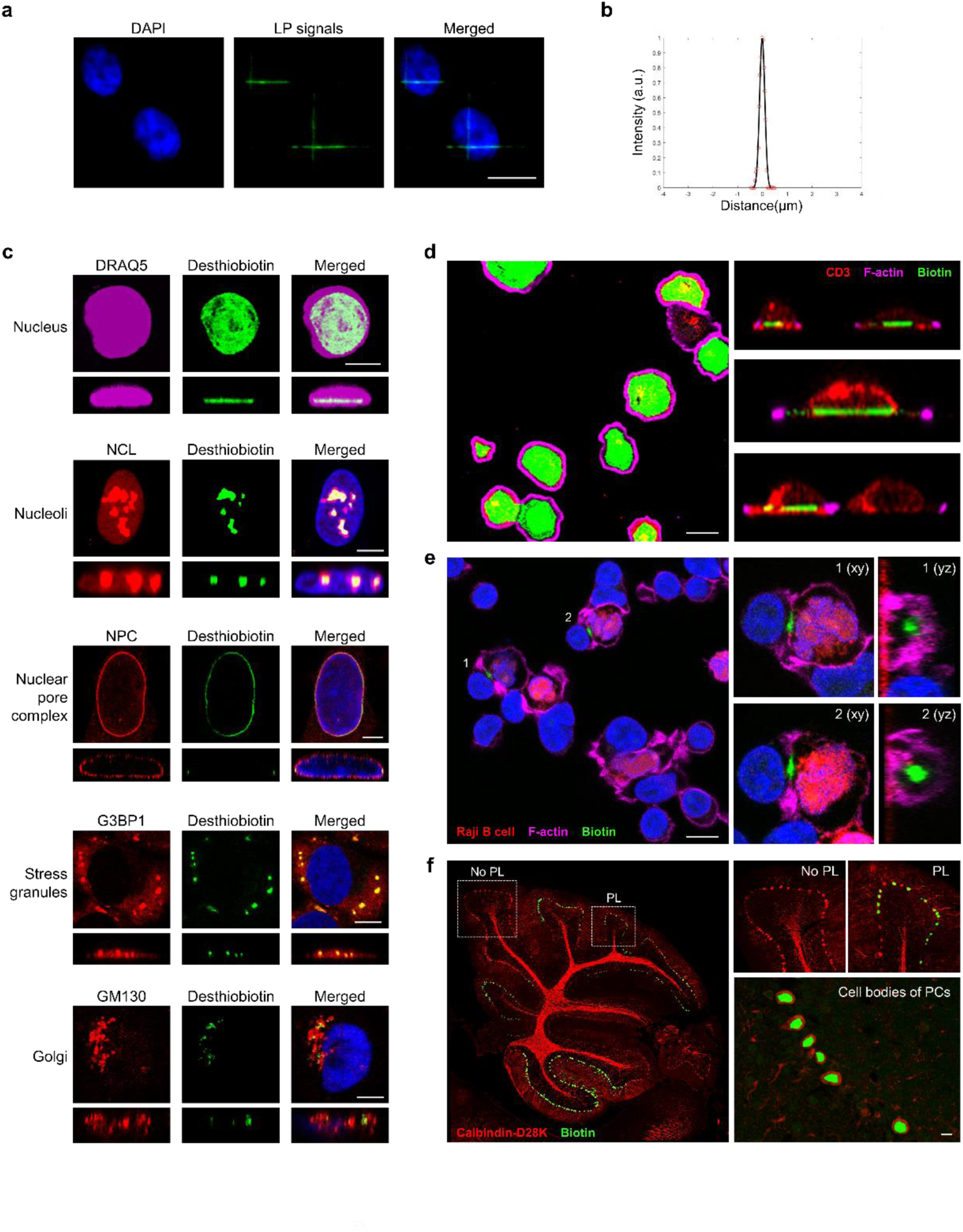
Targeted photo-biotinylation. **a,** A line “cross” pattern is photolabeled on fixed U-2OS cells, and the biotinylated molecules are shown by Dy488-NeutrAvidin. **b,** Photolabeling resolutions when using a 40x/0.95 NA objective, FWHM = 0.24 µm or 240 nm, as shown by super-resolution structured illumination microscopy. **c,** Photo-biotinylation in xy (top view) and z (side view) directions of various subcellular structures, visualized by confocal images. The ROIs are stained with Alexa Fluor 568 secondary antibody, and the photolabeled signals are shown with Dy488-NeutrAvidin. **d,** Precision photo-biotinylation (Dy488-NeutrAvidin) at a thin layer of immune synapses of Jurkat T cells, demonstrated in a spreading assay. **e,** Precision photo-biotinylation (Dy488-NeutrAvidin) of immune synapse of co-cultured Jurkat T cells and Raji B cells, illustrating the capability for two-color image analysis. **f,** Photo-biotinylation (Dy488-NeutrAvidin) of proteins in an FFPE mouse brain tissue section. Cell bodies of Purkinje cells are selected for photolabeling. Scale bar: 10 µm.

### Protein spatial purification enables subcellular spatial proteomics in high sensitivity and specificity

To check whether ultra-content spatial biotinylation yielded isolation of enough proteins for MS sensitivity, we first tested targeted photolabeling and pulldown of nuclear proteins as a proof of principle. Three biological replicates of photolabeling on proteins of fixed U-2OS cells at the regions marked with DRRQ5, a far-red fluorescent DNA intercalating dye, were performed (Fig. 4a). After ∼16-hour targeted photolabeling for each replicate, cells were scraped, harvested, and lysed to extract proteins. Biotinylated proteins were enriched by pulldown using streptavidin beads, tryptically digested on beads, and subject to LC-MS/MS measurement (Fig. 4a). Dot-blot analysis revealed that biotinylated proteins were enriched in the protein lysate and streptavidin beads pulldown when the photolabeling illumination was applied (Fig. 4b), demonstrating effective photo-biotinylation of the sample. Furthermore, the internal control α-tubulin was only present in the lysates and not in the pulldown, indicating the efficiency of the pulldown for nuclear proteins. When BBZP were used for photo-biotinylation, 4,820 proteins were identified with high confidence (Table S1). A distribution of overall protein abundances was binned by the ratio of copies in a photolabeled (PL) sample to those in a control (CTL) sample annotated as PL/CTL ratio (Fig. 4c). The specificity of the spatially labeled nuclear proteome was determined by calculating the percentage of the true positive proteins known as nuclear proteins among the obtained proteome. A total of 1,316 proteins showed differentially enriched, where 1,207 were annotated as nuclear proteins, accounting for a 92% true positive rate. We termed the technique of integrating high-speed ultra-content protein spatial purification and proteomics as optoproteomics.

**Fig. 4.**
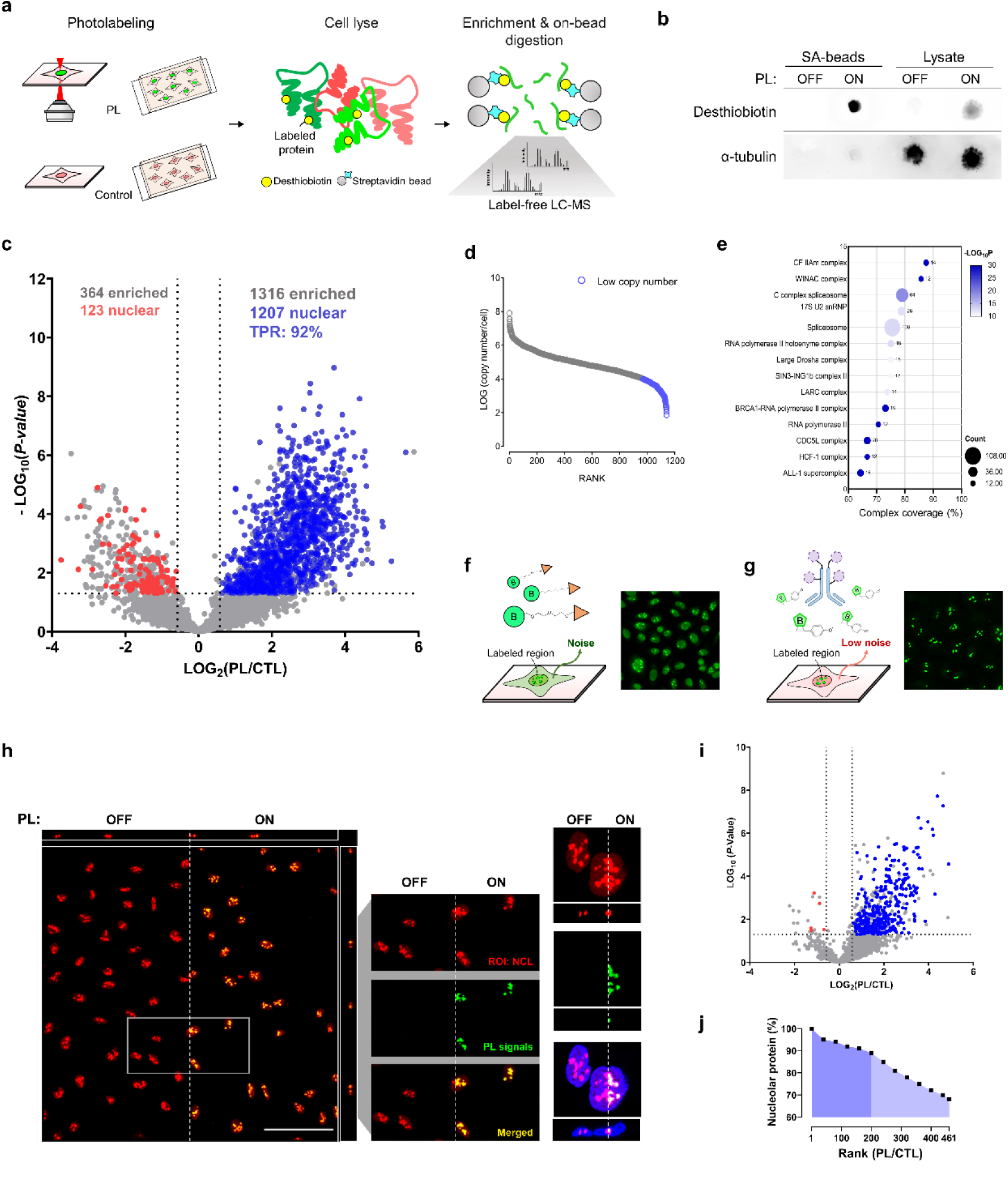
Subcellular proteomics by spatial purification. **a,** Steps of proteomic profiling after targeted photo-biotinylation. Both photolabeled (PL) cells and control cells (without illumination) are lysed, enriched using streptavidin beads, and then digested with trypsin prior to LC-MS/MS measurement. **b,** Effective protein photo-biotinylation as demonstrated by the dot-blot assay, showing desthiobiotin signals in photolabeled (ON) cells but not in control cells (OFF). **c,** High specificity of nuclear proteins obtained through spatial purification with targeted photo-biotinylation. A distribution of overall protein abundances is binned by the ratio of copies in a photolabeled (PL) sample to those in a control (CTL) sample annotated as PL/CTL ratio. High true positive rate of nuclear proteins in the PL enriched group (blue) compared to the CTL sample. **d,** Distribution of protein copy numbers, with the low copy number ones shown in blue (< 10,000 copy number per cell). **e,** CORUM analysis of protein complexes, revealing major nuclear associated complexes. **f-g,** Method comparison using a free photoactive probe (f) and an antibody-conjugated photoactive probe (g). Non-specific biotinylation is reduced when using the antibody-conjugated photoactive probe for photolabeling small compartment (nucleoli: 0.5-3.5 µm). **h,** Precision photo-biotinylation using the antibody-conjugated probe. Confocal images show photolabeled and non-labeled regions of interest (ROIs) within nucleoli, showing precise labeling in xy and z directions using the antibody-conjugated photoactive probe. **i,** Distribution of overall protein abundance according to the PL/CTL ratio for the nucleolar proteome study. **j,** The true positive rate of nucleolar proteins against the rank of PL/CTL for enriched proteins. Scale bar: 10 µm.

Ability to identify low abundant proteins with high sensitivity is important for the discovery of novel biomarkers. We found that more than 10% of the nuclear proteins were identified as low abundant proteins with less than 10,000 copy number per cell (Fig. 4d), illustrating the capability of optoproteomics to identify low abundant proteins. We further examined the capability of protein complex discovery and carried out CORUM complex analysis by employing differentially enriched proteins as seed proteins to reveal protein complexes (Fig. 4e and Table S2) (*31*). The over-represented protein complexes were within our nucleus-targeted region, such as spliceosome, histone complex, and RNA polymerase complex, suggesting optoproteomics not only identified site-specific proteins but also revealed spatially related protein complexes.

### Antibody-conjugated photosensitizers enable high-precision subcellular spatial **proteomics**

To further test the method on smaller subcellular targets, we performed directed photolabeling and following proteomic analysis of nucleoli, the small organelles within the nuclei ranging in size from 0.5 to 3.5 μm. When photolabeling tiny regions like nucleoli, free-diffusing photosensitizers introduced slight expansion of biotinylation areas, reducing the specificity of proteomics (Fig. 4f). To address this issue, we used secondary antibody-conjugated [Ru(bpy)_3_]^2+^ to hybridize the photosensitizer close to the ROIs along with DBP in the media (Fig. 4g). Anti-nucleolin (NCL) primary antibody was employed as the nucleolar marker. Images were analyzed to segment the nucleoli from the nuclei to avoid undesired labeling of antibodies outside of the nucleoli. High-precision biotinylation at the nucleoli was achieved using microscopy-guided photolabeling (Fig. 4h). Labeling background even in the highly condensed nuclei was low, illustrating that the method of using antibody-conjugated photosensitizers can be broadly applicable to subcellular affinity labeling. Our proteomics results yielded 3,162 proteins with high confidence (Table S3). The distribution of protein abundances was plotted by comparing the ratio of photolabeled nucleoli (PL) against the control sample (CTL) (Fig. 4i). The specificity of the spatially labeled nucleolar proteome was determined by ranking the abundance based on the PL/CTL ratio. Notably, within the top 200 ranked proteins, 180 were annotated as nucleolar proteins, resulting in a true positive rate of 90% (Fig. 4j).

### Optoproteomics enables the discovery of novel protein constituents of arsenite-induced stress granules

Beyond validation of known proteomes, the key question is whether optoproteomics has a discovery capability. Stress granules (SGs) are dynamic ribonucleoprotein assemblies that are formed under cellular stress (*32*). Given their pathological implications in neoplastic and neurodegenerative processes, characterization of SG composition is crucial for therapeutic explorations (*33*). The existing proteome was obtained from biochemical fractionation and proximity labeling (*34*), but whether it is extensive enough remains to be checked due to the diminutive, membrane-less structure of SGs. Here, SGs were induced in U2-OS cells via arsenite exposure and stained with SG marker G3BP1. Due to the small size of SGs (∼ 200 nm for small ones) (*35*), we used antibody-conjugated [Ru(bpy)_3_]^2+^ hybridized to the G3BP1 primary antibody for confined spatial photolabeling (Fig. 5a and Fig. S6). In total, 2,785 proteins were identified with high confidence. With a high correlation across replicates, 1,754 intersected proteins of three biological replicates were found (Fig. 5b and Table S4). Applying the log_2_ fold-changed cutoff of 0.585, the number of unique peptides of 3, and the score sequest HT of 100 as the selection criteria, we retained 124 significantly enriched proteins (Fig. 5c). Many well-known SG proteins including hnRNPs, eRF3a, PABP1, TADBP, FXR1, and eIF3s were enriched with high PL/CTL ratios. Gene ontology enrichment analysis showed that the identified proteome was highly associated with cellular macromolecule biosynthetic process, cellular component biogenesis and organization, RNA and nucleobase-containing compound metabolic process, and cellular protein localization (Table S5). Surprisingly, 40% of these 124 proteins were absent in the existing SG proteome. These enriched proteins without prior annotation support as SG proteins exhibited high interaction relationship with the SG proteins in the association network (Fig. S7).

**Fig. 5.**
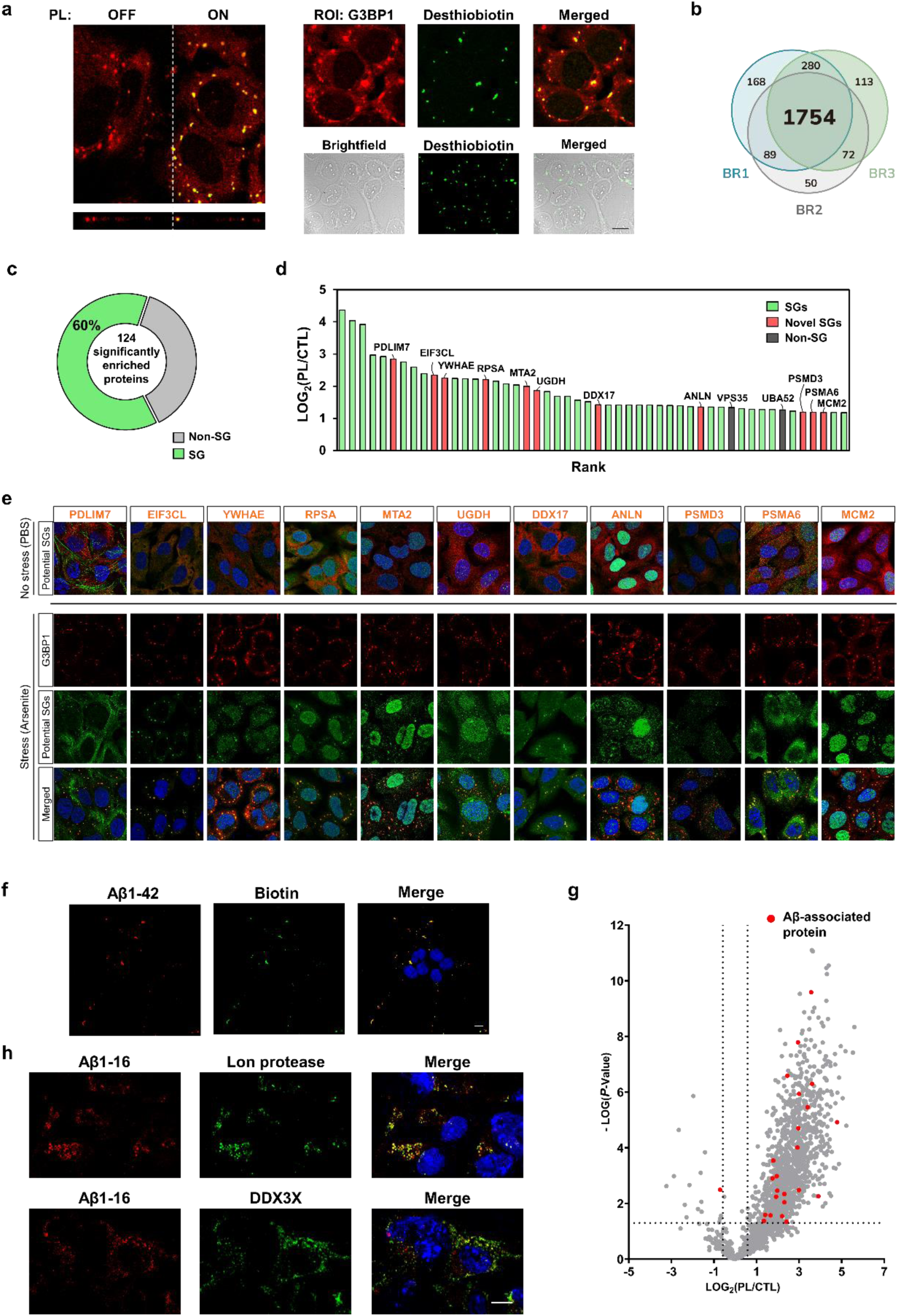
Novel protein constituents discovered by optoproteomics. **a,** Confocal images depicting precise and accurate photolabeled stress granules (SGs) by the antibody-conjugated ruthenium probe. **b,** Venn diagram of three biological replicates of spatial photolabeling SG proteomics. **c,** The percentage of true positives when comparing the significantly enriched proteins from spatial purification versus the SG proteins in the existing database. **d,** Identification of potentially novel SG protein constituents among top-ranked spatially purified proteins using optoproteomics. Known SG proteins are shown in green, whereas positively validated ones (see below) are shown in red. The ones not validated are shown in gray. **e,** Validation of a series of novel SG proteins by immunostaining. SG formation of U-2OS cells is stimulated by arsenite stress. Proteins identified as potential SG proteins (green) are highly co-localized with SG marker G3BP1 (red). **f,** Confocal images depicting precise and accurate photolabeling of amyloid-β (Aβ) plagues using the BBZP probe. **g,** Distribution of overall protein abundance according to the photolabeled/control (PL/CTL) ratio for the amyloid-β plague proteome study. **h,** Two highly ranked proteins (Lon protease and DDX3X) not previously associated with Aβ aggregates were found to colocalize with Aβ in the differentiated SH-SY5Y cells. Scale bar: 10 µm.

To examine whether the specificity of the proteomics result was not good enough or we revealed novel SG-associated proteins, we used immunostaining to test colocalization of G3BP1 and each of the proteins without prior annotation support as SG proteins at a time (Fig. 5d). Among 13 proteins tested, 11 of them were colocalized with G3BP1, i.e., PDLIM7, EIF3CL, YWHAE, RPSA, MTA2, UGDH, DDX17, ANLN, PSMD3, PSMA6, and MCM2 (Fig. 5e). Considering all known SG proteins and these validated SG-localized proteins, the SG specificity of the obtained proteome reached 96% among the top 50 identified proteins, demonstrating the discovery power of optoproteomics with its high specificity. Notably, the role of PDLIM7, DDX17, and PSMA6 in cell differentiation illustrated a potential crosstalk between stress response and cell development, as their arsenite-triggered cytosolic localization suggests a pivot from normal cellular activity to stress management, contributing to the SG formation.

### Optoproteomics enables discovery of novel constituents in amyloid-β

To further examine optoproteomics usage on disease-related problems, we applied it to amyloid-β (Aβ), peptides aggregating into insoluble plaques that characterize the Alzheimer’s disease (*36*). Targeted protein photo-biotinylation was performed at the locations of Aβ1-42 aggregates overexpressed in differentiated human SH-SY5Y neuron-like cells, a cell model for neurodegenerative diseases (Fig. 5f). Applying optoproteomics, we identified a total of 1,378 proteins associated with Aβ1-42 (Table S6), inclusive of 23 established markers intrinsic to Aβ peptide aggregates (Fig. 5g). Intriguingly, bioinformatic analysis unveiled a consistent enrichment of proteins tied to key neurodegenerative diseases like Alzheimer’s, Huntington’s, Parkinson’s, and amyotrophic lateral sclerosis across three biological replicates, with a notable focus on acetylation and phosphorylation processes (Table S7) (*37, 38*). Among the high-ranked enriched proteins, several candidates not previously directly associated with Aβ aggregates were identified. Two of these candidates, DDX3X, a DEAD-box RNA helicase, and Lon protease, a mitochondrial serine peptidase, were found to colocalize with Aβ1-16 as shown by immunostaining, suggesting their potential association with amyloid-β (Fig. 5h). Aggregation of DDX3X in amyloid β may trigger inflammatory cell death (*39*), whereas accumulation of Lon protease in amyloid β may impair mitochondrial function and cause cell death (*40*). Although it is unclear whether these two proteins a play substantial role in Alzheimer’s disease, optoproteomics was able to reveal novel amyloid-β-localized proteins in a cell model for further functional studies.

## Discussion

Spatial proteomic discovery in submicron specificity and low-abundance sensitivity is the unique strength of optoproteomics. With the platform’s ultra-content targeted photo-biotinylation capability, identifying novel protein constituents at a specific subcellular location under a microscope from cell or tissue samples becomes feasible, giving scientists an unprecedented tool to largely and precisely expand the protein “proximome” understanding for specified ROIs. The application scope of optoproteomics is broad. For example, one can determine proteins at the tip of primary cilia, cytosolic TDP-43 aggregates, kinetochore microtubules, PD-L1 adjacent sites of SOX2-enriched cells of a tumor tissue, the lipid droplet-mitochondria interacting region of ballooning hepatocytes from a non-alcoholic steatohepatitis (NASH) patient sample, to name a few. Although the finding is a proximome instead of a direct interactome, the high specificity somewhat guarantees the appearance of identified proteins in the neighboring region of an ROI, providing a robust list of proteins for hypothesis testing and following mechanism investigations. Thus, optoproteomics is expected to largely increase the total location-specific proteome database useful for revealing potential disease-associated protein players and providing testable hypotheses related to protein interaction partners for basic research. Currently the major alternatives of unbiased high-precision proteomics are proximity labeling including the use of APEX2, BioID, and TurboID (*10, 12, 41*), as well as LCM, especially the recent AI-enabled DVP dissection (*21*). For problems where a highly specific bait can be used, proximity labeling has a strong advantage over optoproteomics in sensitivity, capable of pooling from multiple 10-cm dishes to collect enough proteins for MS analysis. It surprised us that the study of SGs described above was able to identify tens of novel constituents compared to the existing SG proteome database that incorporates results from a previous proximity labeling study, potentially due to the purity of protein collection through microscopy-guided spatial filtering. Another advantage of optoproteomics is to study human FFPE tissue samples, where no exogenous expression is needed. Regarding the comparison with LCM, if the area of interest is large such as a multicellular region of a cancer tissue sample, DVP-based LCM should be the method of choice, because it has a much better throughput than optoproteomics. Furthermore, LCM does not require the pulldown step, largely reducing the sample loss through the enrichment process. When the targeted region is in the submicron range, even though the LCM can aim the targets in high precision, the required laser burning energy produces a varying laser beam width of 2-4 µm depending on the laser moving speed, making it relatively challenging to dissect small elements. Another difference between LCM and optoproteomics is that the LCM collects samples from the entire dissecting column, whereas optoproteomics specifies the axial location of the sample through either two-photon focusing or antibody-conjugated confined reactions, excluding non-specific proteins from undesired planes.

Optoproteomics supplements existing technologies in spatial biology with its capability of unbiased subcellular spatial proteomics discovery. Several new methods have recently been developed for high-precision spatial biology, specifically for targeted/*de novo* spatial transcriptomics and targeted spatial proteomics (*1–4, 42–44*). When aiming to understand cell heterogeneity of a mm-scale tissue, several existing spatial biology technologies are valuable and have advantageous throughputs to generate an intercellular omics mapping. If a biological problem is to understand the molecular mechanisms or the drug mechanisms of action at the protein level, one would prefer to validate spatial transcriptomics results with further protein studies. When subcellular localizations are crucial for understanding protein interactions, locations of corresponding mRNAs and proteins may differ (*45*), and thus subcellular proteomic discovery using a method such as optoproteomics becomes uniquely valuable. Once a novel proteome is discovered by optoproteomics, existing antibody-based spatial proteomics approaches will be useful for large-scale localization validation.

Speed optimization enables the optoproteomics technology to perform subcellular spatial discovery within a manageable time. Previously it would take years of work to identify >10 high-specificity protein constituents at a ROI such as β amyloids from Alzheimer’s patient samples. Now with optoproteomics, one can identify a series of novel protein constituents specifically localized at β amyloids through three or more repeats of rounds in about 2-3 months (including potential AI development), and then bought the corresponding antibodies to perform colocalization validation within another couple of months, largely increasing the throughputs of future disease-associated protein identification.

There are rooms to improve the photolabeling speed of optoproteomics, because ∼10 hours for labeling 5 × 10^5^ cells still appear to be long. One way to accelerate the process is to use a widefield illumination instead of a point scanning one. It is possible to increase the peak power of a femtosecond laser by using a lower repetition rate to satisfy the energy requirement for widefield two-photon temporal focusing (*46, 47*), but the setting will be more involved. If a single-photon light source is used instead of a two-photo one, and with a widefield pattern illumination setup such as a digital micromirror device, the process speed can also be expedited. However, scattering due to heterogeneous refractive index of a biological sample remains to be a challenge for high spatial precision photochemistry using a single-photon light source. Another possibility is to improve the efficiency of photochemical reaction so that one can improve the labeled protein copies with the same reaction illumination duration. Development of new or modified photochemical probes will be needed to reach this goal. Yet another possibility to accelerate the process is to push the sensitivity limit toward single cell proteomics, including the use of nanoPOTS platform or the recent advanced mass spectrometers for single cell proteomics (*48–50*). Although the goals of single cell proteomics and subcellular proteomics are different, where single cell proteomics aims to understand cell heterogeneity and subcellular proteomics aims to discover location-specific protein constituents, the technology advent for single cell proteomics does help improve the sensitivity of subcellular proteomics using a method like optoproteomics.

In addition to photo-induced protein biotinylation, there are choices of tags other than biotin, such as SNAP-tag, CLIP-tag, HaloTag, or click chemistry. The photoreactive warheads also have different choices, including diazirine, phenyl azide, and CNVK for RNA or DNA. Warheads reactive to different wavelengths are also possible. With a combination of different tag and warhead choices, it is possible that the optoproteomics method can be extended to spatial multi-omics studies such as for two or more species of proteome targets or proteome-transcriptome targets for a more comprehensive omics understanding of a biological problem.

## Supporting information

Photolabeling video 1

Photolabeling video 2

Photolabeling video 3

## Acknowledgements

We thank Shean-Jen Chen, Chia-Yuan Chang, Huan-Cheng Chang, and Hsiao-Hua Yu for advice and help. We thank the staff of the Medicinal Chemistry and Analytical Core Facilities at National Biotechnology Research Park supported by Academia Sinica AS-NBRPCF-111-201 during this project. This work was supported by Academia Sinica Taiwan Instrumentation Development Program, Academia Sinica AS-SUMMIT-108, Ministry of Science and Technology, Taiwan 108-2823-8-001-002, and Syncell Inc.

