## Supplementary information for "Microscopy-guided subcellular proteomic discovery by high-speed ultra-content photo-biotinylation"

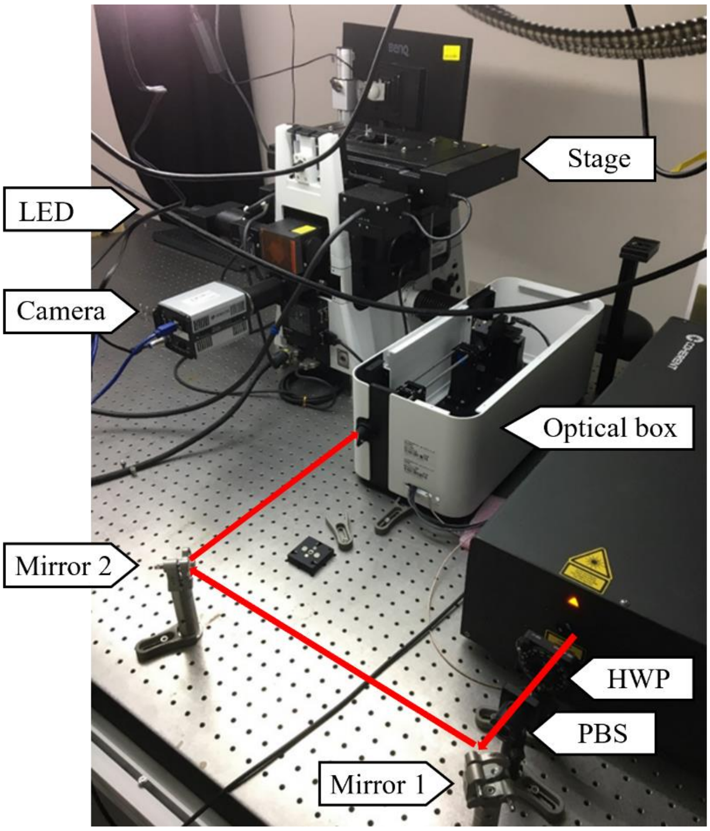


**Figure S1**.

An image-guided two-photon illumination microscope system (back view). The red arrows indicate the path of the input laser beam for the optical box containing a pair of galvanometers and other optical components. This optical box is connected to the side port of an inverted microscope, which is mounted with a camera and an LED imaging light source.

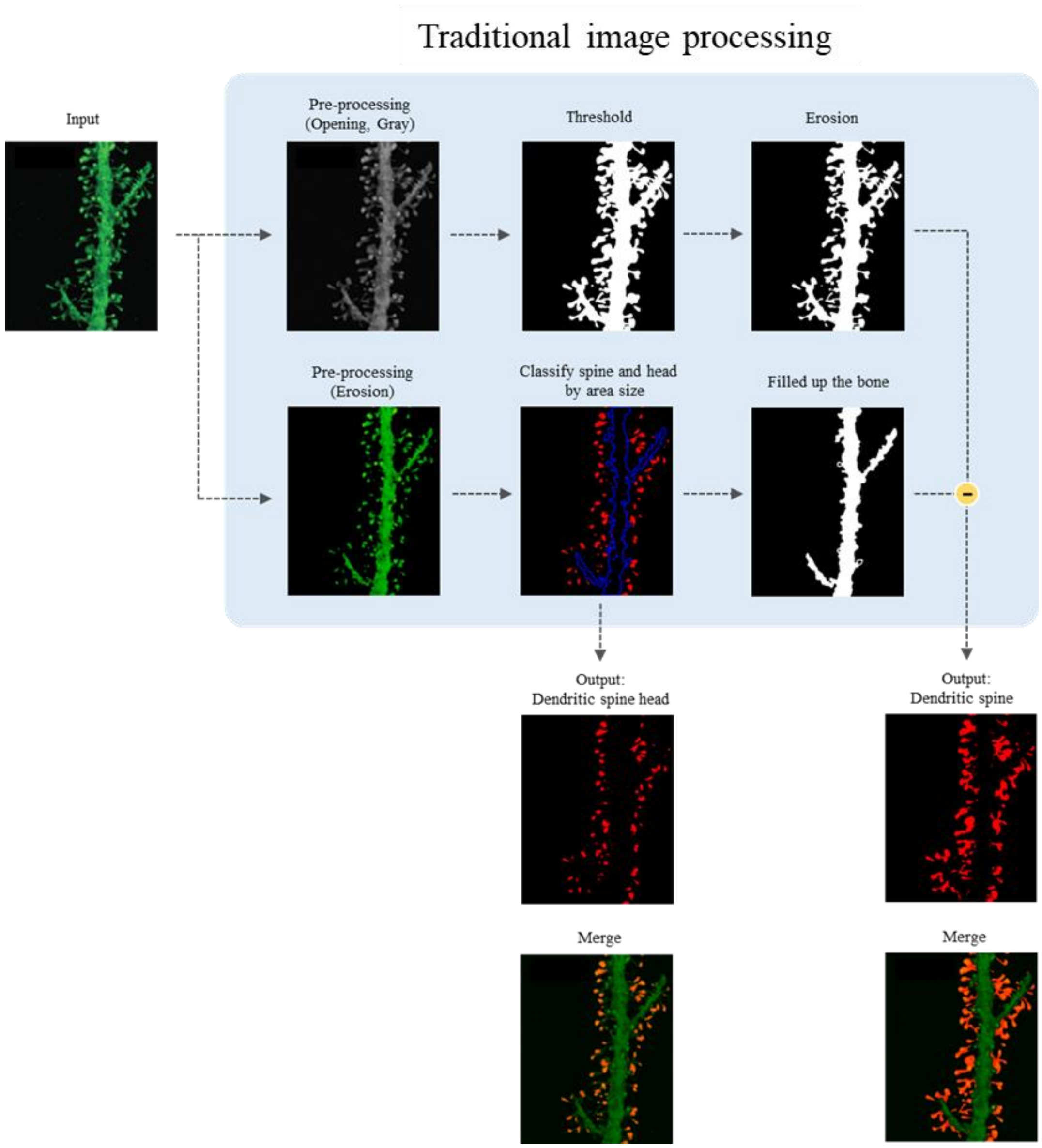


**Figure S2.**

An example of image segmentation of the dendritic spine heads and the dendritic spines through a series of traditional image processing steps.

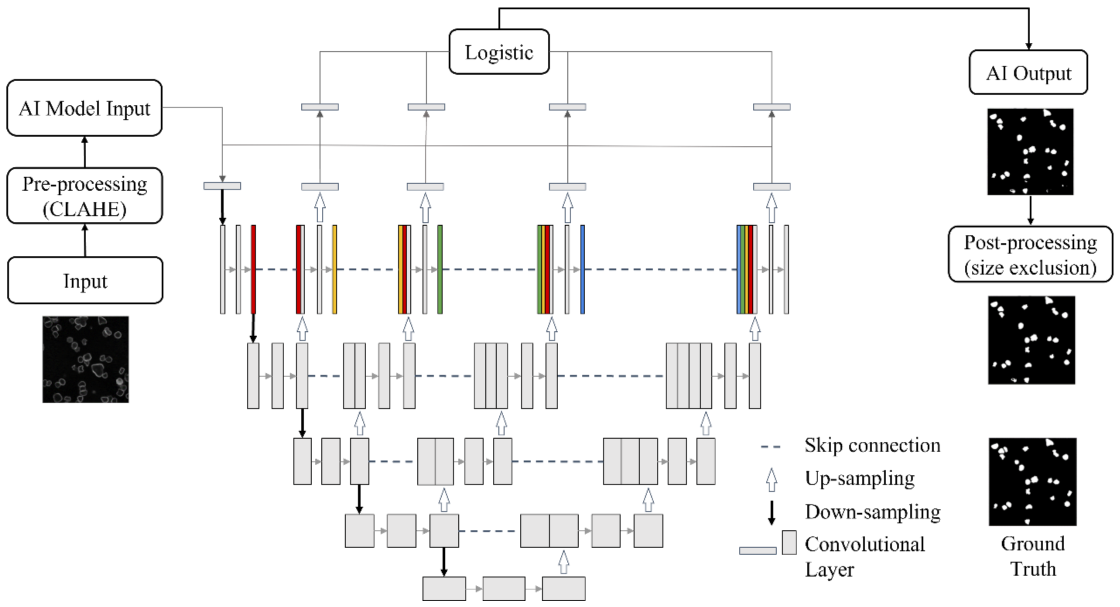


**Figure S3.**

An example of using a trained AI model to segment active immune synapses of a spreading assay sample. A U-Net model is used for image segmentation. A post-processing is applied on the AI output to select the regions within a desired range of sizes. The output result shows high accurate quality, which can be validated by the ground truth image (annotated result). This image segmentation process takes only about 200 ms for each field of view.

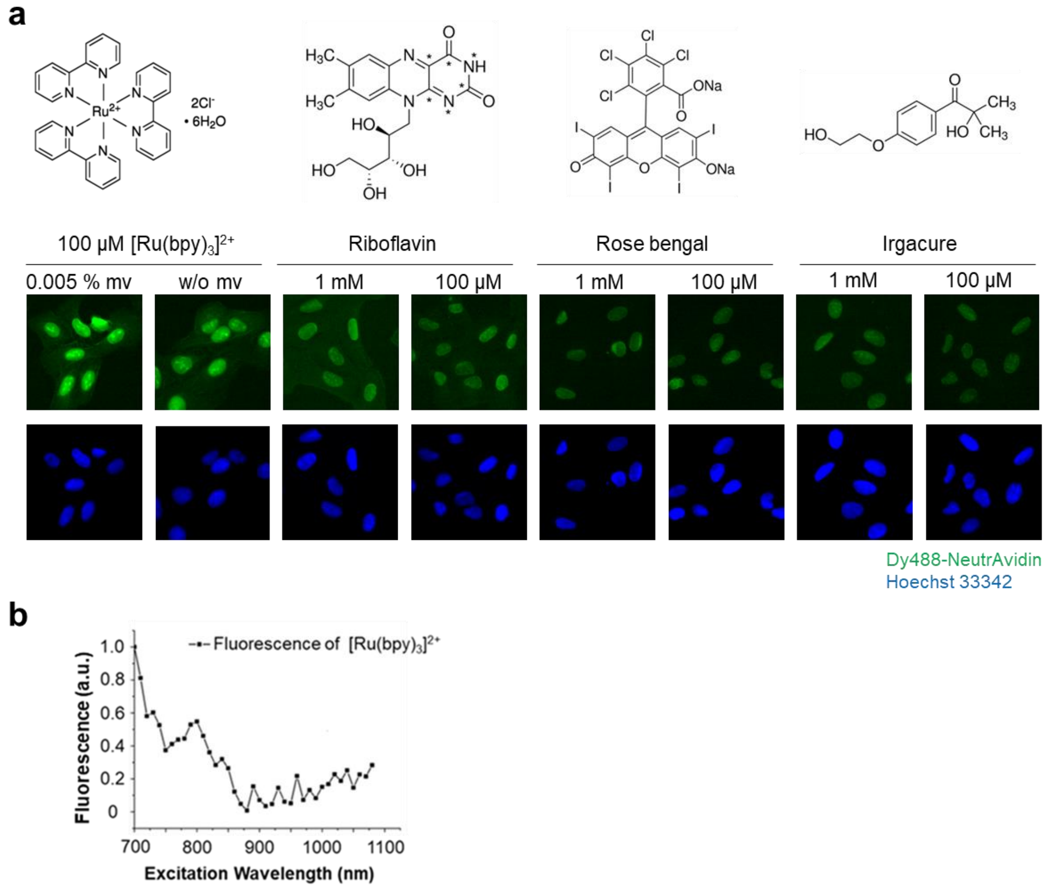


**Figure S4.**

**a)** Four commercially available photoactivate probes: $[Ru(bpy)_3]^{2+}$, riboflavin, rose bengal, and Irgacure (upper panel). Photolabeling results of U-2OS cells immersed with desthiobiotin-phenol and illuminated with two-photon illumination scanning through the entire field of view at wavelength ( λ ) = 780 nm and power = 100 mW. Highest fluorescent signals are detected when using $Ru[(bpy)_3]^{2+}$ with 0.005% methyl viologen (mv).

**b)** Fluorescence emission spectrum of $[Ru(bpy)_3]^{2+}$ excited by femtosecond laser illumination between 750 - 820 nm.

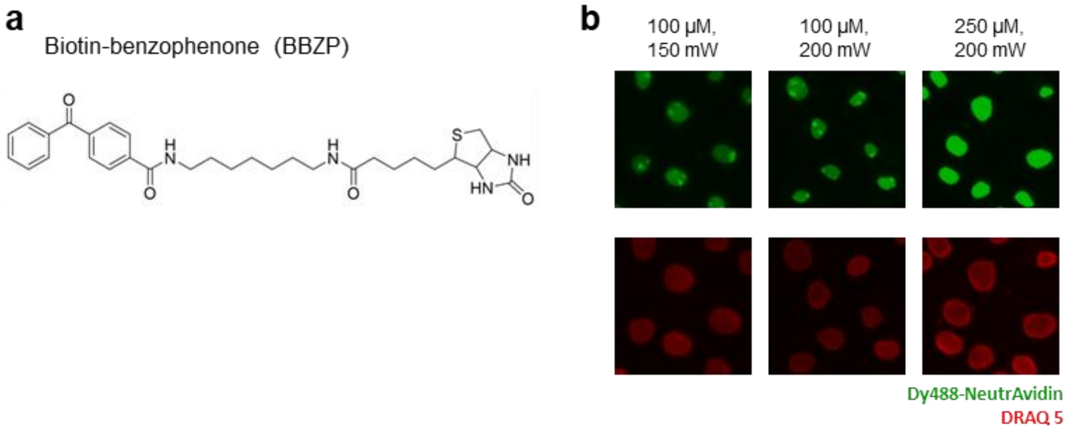


**Figure S5.**

**a)** Illustrated structure of biotin-benzophenone (BBZP). **b)** Biotin labeling efficiency is increased with the labeling laser power and the concentration of BBZP.

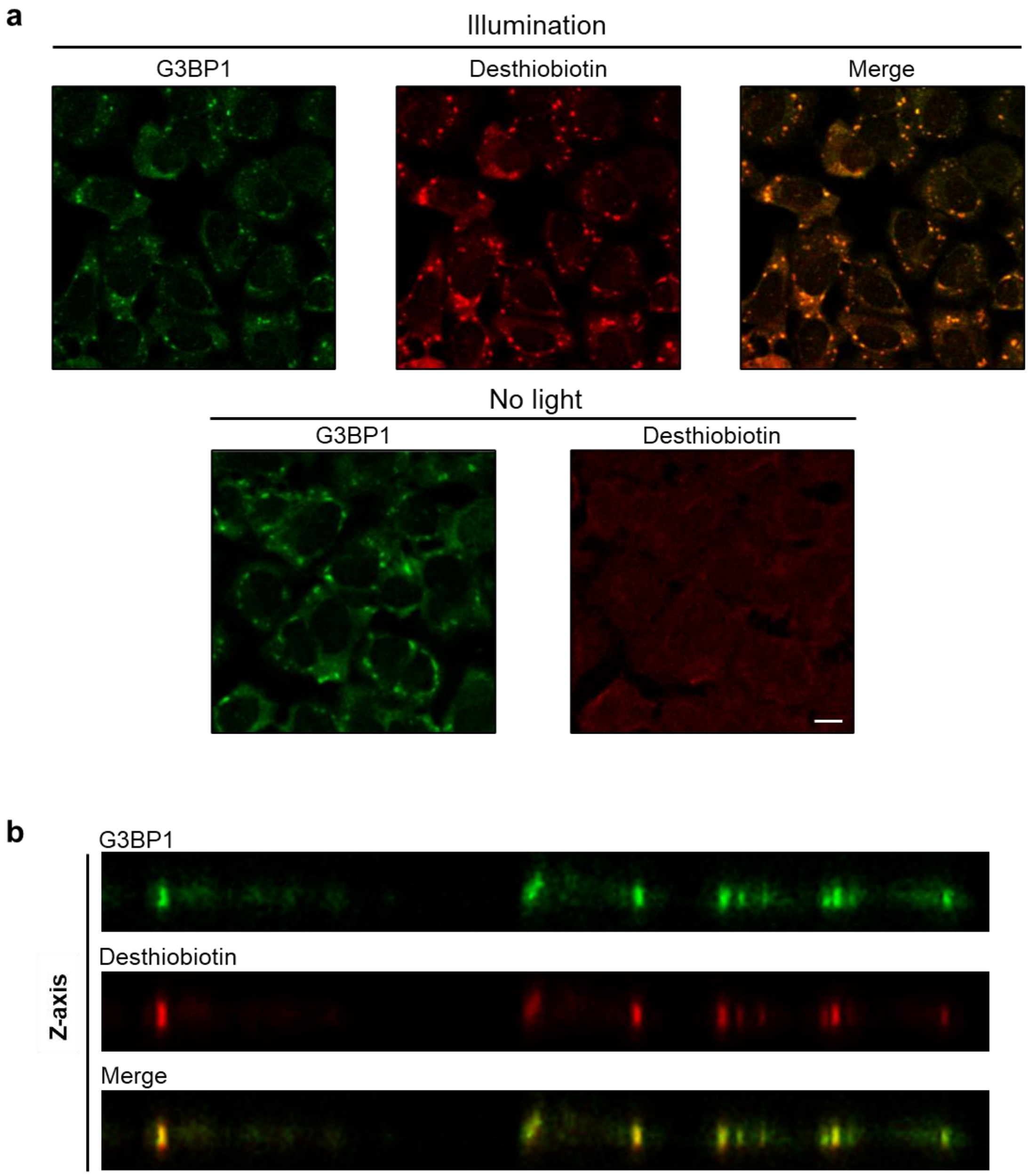


**Figure S6.**

**a)** Precision labeling of stress granules of U-2OS cells induced by arsenite treatment using targeted photo-biotinylation by two-photon illumination. Labeling signal is negligible when two-photon illumination is absent. Scale bar: 10 μm.

**b)** Precision labeling in the axial direction.

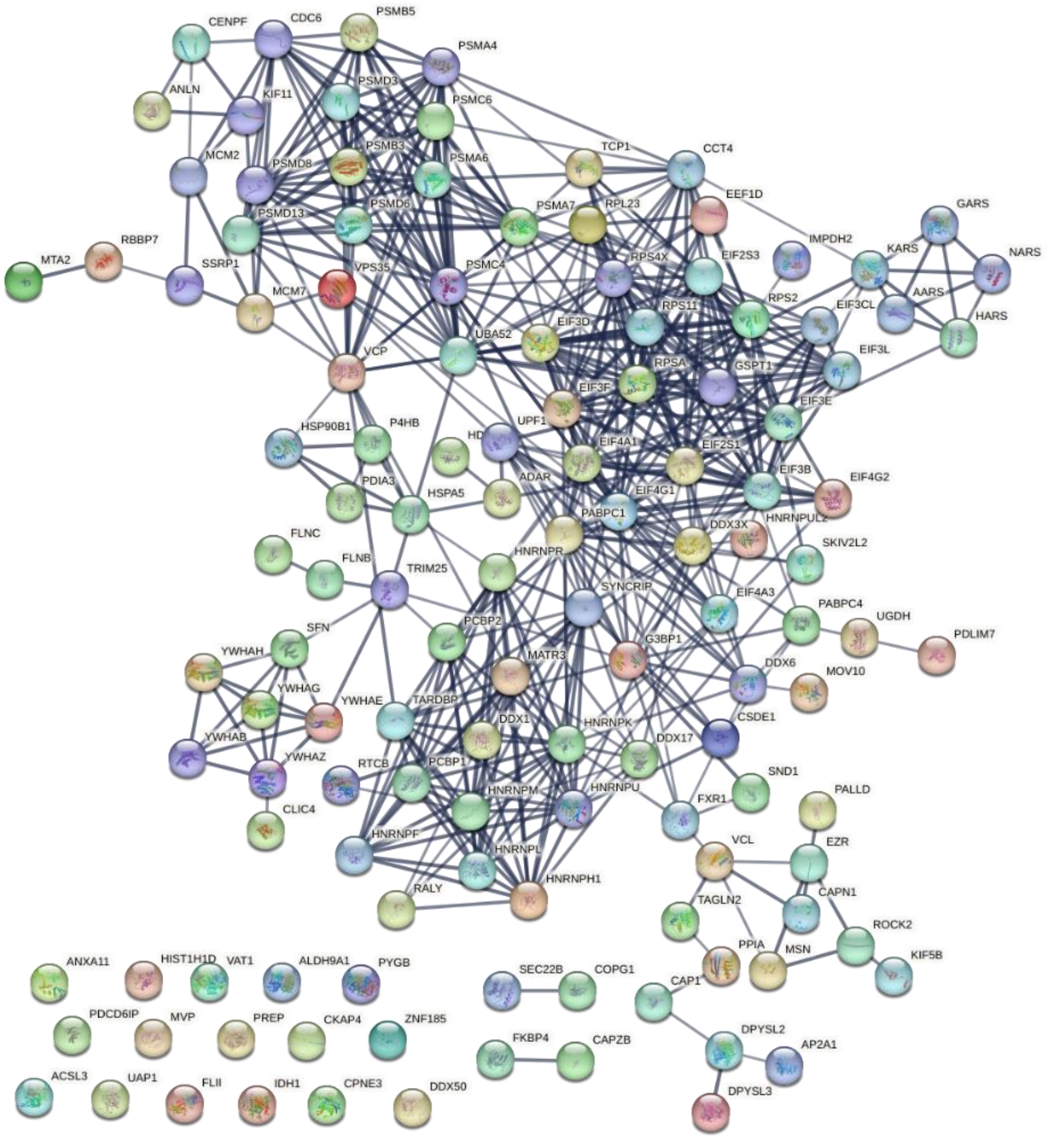

**Figure S7.**
STRING analysis of the stress granule proteome obtained by optoproteomics. The enriched 124 proteins are subject to STRING analysis to reveal functional and physical interactions among the proteins. Each node represents a protein, and each edge represents a degree of physical and functional interactions.
