## Supplementary material for "Microscopy-guided subcellular proteomic discovery by high-speed ultra-content photo-biotinylation": Materials and methods

Cell preparation

U-2OS cells (HTB-96, ATCC, VA, USA) were cultivated at 37°C in a 5% $CO_2$ humidified environment in Dulbecco's Modified Eagle Medium (DMEM, 11965092, Thermo Fisher Scientific, MA, USA) supplemented with 10% FBS. $2 \times 10^5$ cells were seeded in glass bottom chambers (80287, ibidi, WI, USA) and incubated for approximately 16 h to reach 80% of confluency. Afterwards, cells were washed with phosphate buffered saline (PBS) and then fixed with 2.4% paraformaldehyde solution (PFA, 15710, Electron Microscopy Sciences, PA, USA). For the induction of stress granules, U-2OS cells were similarly cultivated and seeded in glass bottom chambers and incubated for approximately 16 h to reach 90% of confluency. These cells were then treated with 500-μM sodium arsenite solution (S7400, Sigma-Aldrich, MO, USA) for 1 h at 37˚C in the incubator, and fixed with 100% ice-cold methanol. Fixed samples were incubated with PBS/0.5% Triton X-100 to permeabilize the cell membrane and blocked with 3% bovine serum albumin (BSA) in PBS/0.1% Triton X-100 (PBS-T) for 1 h, followed by 30 min of 0.002% streptavidin and 15 min of 40 μM biotin blocking.

Spreading assay of Jurkat T cells

Glass chambers were coated with poly-D-lysine (P7280, Sigma-Aldrich), washed with PBS and incubated with 10 mg/mL mouse anti-CD3 (317301, BioLegend, CA, USA) and 10 mg/mL anti-CD28 (302933, BioLegend) antibodies in PBS overnight at 4˚C. Jurkat T cells (Clone E6-1, TIB-152 , ATCC) were dropped on the glass chamber and incubated at 37 ˚C. After 15 min of incubation, the Jurkat T cells were fixed with 4% PFA. To visualize the T-cell synaptic contact, immunostaining of CD3 with anti-mouse AlexaFluro 568-IgG (A-11004, Thermo Fisher Scientific) as the secondary antibody and phalloidin-Atto 647 (65906, Sigma-Aldrich) was performed.

Immune synapse induction

Raji B cells (CCL-86, ATCC) were utilized as antigen presenting cells (APCs). To activate these APCs for immune synapse (IS) formation, $1 \times 10^6$ Raji B cells were pelleted to resuspend with 100 µl serum-free RPMI (SH30027, HyClone, UT, USA) containing 0.1 µg Staphylococcal Enterotoxins (ET404, Toxin Technology, FL, USA) and incubated at 37°C for 1 h. On the last 20 min of the incubation, the Raji B cells were stained with 20 µM cell tracker Red CMTPX Dye (C34552, Thermo Fisher Scientific). The cells were then washed twice with PBS to remove excess dyes. $6.5 \times 10^5$ Raji B cells were then resuspended with 500-µl serum-free RPMI and seeded on individual well of a poly-D-lysine coated glass chamber for 20 min. Equal amount of Jurkat T cells were resuspended with 250-µl serum-free RPMI and then added dropwise into the well containing the Raji B cells. After a 10-minute incubation at 37°C, the T-B cell conjugates were fixed with 2% PFA solution for 10 min, followed by 100% ice-cold methanol fixation for 5 min at -20°C. To visualize T-B cell IS, staining of CellTracker Red CMTPX dye (C34552, Thermo Fisher Scientific) and phalloidin-Atto 647N was performed.

Amyloid sample preparation

SH-SY5Y cells (CRL-2266, ATCC) were cultivated at 37°C in a 5 % CO2 humidified environment in DMEM supplemented with 10% FBS. To induce differentiation, the SH-SY5Y cells were cultured in serum-free neurobasal medium (21103049, Life Technologies, NY, USA) for 5 days, and then infected with lentivirus to express mCherry-Aβ1-42. The formation of Aβ1-42 aggregates was facilitated by maintaining the cells in culture for 72 h post-lentivirus infection and then an additional 48 h to allow for aggregate formation. The SH-SY5Y cells were fixed by 2.4% PFA.

Tissue sample preparation

A C57BL/B6 mouse (Jackson Laboratory, ME, USA) was perfused with 0.9% NaCl to clear the blood, followed by fixation with 4% PFA. The mouse brain was processed using the formalin-fixed paraffin-embedded (FFPE) block procedure, which included fixation, dehydration, and paraffin embedding. FFPE slices were sectioned at a thickness of 10 µm, then deparaffinized and rehydrated. Purkinje cells were immunostained with anti-calbindin-D28K antibody (C9848, Sigma-Aldrich) overnight at 4°C, followed by immunostaining with the secondary antibody, AlexaFluor 555 (A-32727, Thermo Fisher Scientific).

Optical setup

A femtosecond laser (Chameleon Vision I: Coherent, CA, USA) was used for two-photon illumination at 140-fs pulse, 80-MHz repetition rate, 780-nm center wavelength, and -24,000 $fs^2$ pre-compensation. The laser power was adjusted manually by rotating a half-wave plate (HWP) together with a polarizing beam splitter (PBS). The on/off switching of the laser was modulated by an acousto-optic modulator (AOM) (AOMO 3080-125: Gooch & Housego, UK). A quarter wave plate (QWP) was used to form circular polarization to achieve homogeneous two-photon labeling within a field of view (FOV). A pair of lenses were used between the AOM and the QWP for beam expansion. The laser beam was steered by a pair of galvanometers (6215H scanners with 671 drivers: Cambridge Technology, now Novanta, NC, USA) and passed through an inverted microscope (Eclipse Ti2-E: Nikon, Japan) containing a scan lens (SL), a tube lens (TL), a long-pass dichroic mirror (DM2) (FF765-Di01: Semrock, NY, USA) at the bottom filter cube,  a multi-band dichroic mirror at the top filter cube (DM1) (FF408/504/581/667/762-Di01: Semrock), and then focused onto the sample plane by an objective of the microscope (CFI Plan Apochromat Lambda 40X, N.A. 0.95: Nikon). The two-photon labeling was guided by images from fluorescence imaging with an LED illumination source (pE-4000: CoolLED, UK) installed with excitation filters for each channel (FF02-482/18, FF01-560/14, FF01-642/10: Semrock). Imaging was captured using an sCOMS camera (Zyla 4.2P USB3: Andor, Oxford Instruments, UK), paired with a multi-band emission filter (FF01-440/521/607/700: Semrock) placed in the bottom filter cube.

Electronics control

A desktop computer was equipped with a field programmable gate array (FPGA) module (NI PCIe-7857: National Instruments, TX, USA) through a PCIe interface. The FPGA module offered multifunction reconfigurable I/O, which allowed digital control of the galvanometers and transistor-transistor logic (TTL) control of the AOM for modulating the femtosecond laser. A main program managed the FPGA and control of other hardware, including the sCMOS, the LED, the microscope stage, and the microscope’s focus drift-free system (PFS: Nikon). Coordinates of a calculated scanning path were encoded and sent to the FPGA to control the galvanometers and the AOM.

Image segmentation by traditional image processing

Different image processing techniques and combinations, including thresholding, erosion, equalization, and filtering, were employed for image segmentation of different patterns of interest. Specifically, for primary cilia, due to their small size, the morphological tophat method was used to extract the target while avoiding extraction of excessively large areas. Basic thresholding was then used to extract the cilia. For nuclear membrane, contrast-limited adaptive histogram equalization and a Gaussian image filter were used as image preprocessing steps. Basic thresholding was then applied for object (nucleus) extraction. To define the edge of each object, the binary result of the object was subtracted from the binary result with light erosion.

Image segmentation by deep learning

Convolution neural network (CNN) was modified from U-Net using pre-trained weights from ImageNet to facilitate convergence for deep learning semantic segmentation. The RMSprop optimizer was utilized to converge the dice coefficient loss function. Following multiple training epochs, the model's stability and accuracy were assessed using a modified metric. Contrast limited adaptive histogram equalization was applied for image preprocessing. For post-processing of the model's output images, erosion was applied to the pattern to minimize the laser intensity's effect on the target. This trained model was applied identically to each FOV.

Conjugation and hybridization of the ruthenium-based antibody

$Ru(bpy)_3$ NHS-ester (Bis(2,2'-bipyridine)-4'-methyl-4-carboxybipyridine-ruthenium N-succinimidyl ester-bis(hexafluorophosphate), 96632, Sigma-Aldrich) was conjugated to the donkey anti-rabbit/donkey anti-mouse IgG antibody (711-005-152/715-005-150, Jackson ImmunoResearch, PA, USA) via an amide coupling reaction. The antibody was reacted with $Ru(bpy)_3$ NHS-ester in a ratio of 0.5 mg/mL antibody to 0.35 mM of $Ru(bpy)_3$ NHS-ester in 100 mM borate buffer (pH 8.0). The reaction was performed at room temperature (RT) for 2 h in the dark. Glycine was added to a final concentration of 100 mM to inactivate the reaction, and the antibody-$[Ru(bpy)_3]^{2+}$ conjugates were then purified using an off-line size exclusion column (PD midiTrap G-25, Cytiva, UK) under a gravity flow.

For hybridization, cells were first incubated with primary antibodies (rabbit anti-NCL (ab22758, Abcam, UK), mouse anti-G3BP1 (sc-365338, Santa Cruz, TX, USA), mouse anti-GM130 (ab52649, Abcam), mouse anti-NPC (ab24609, Abcam)) in PBS-T with 3% BSA for 2 h at RT. After washing with PBS-T, 10 μg/mL of antibody-$[Ru(bpy)_3]^{2+}$ conjugates were hybridized to the primary antibody overnight at 4°C. Cells were immunostained with fluorescence marker (goat anti rabbit AlexaFluor 647: A21245, Thermo Fisher Scientific, or goat anti mouse AlexaFluor 647: A-21235, Thermo Fisher Scientific) for 1 h at RT.

<u>Photolabeling</u>

Cells were incubated with photolabeling reagent containing 5-7 mM desthiobiotin-phenol (LS-1660, Iris Biotech, Germany) and 0.005% methyl viologen (856177, Sigma-Aldrich), and $[Ru(bpy)_3]^{2+}$ (544981, Sigma-Aldrich), Riboflavin (R4500, Sigma-Aldrich), R ose bengal (330000, Sigma-Aldrich), Irgacure (410896, Sigma-Aldrich), or BBZP (Development Center for Biotechnology, Taiwan). Photolabeling was performed using a two-photon laser coupled with a microscopic system at a laser power of 100-200 mW and the cells were subject to a laser exposure time at 100-500 μs. Labeled cells were immediately quenched with the buffer containing 10 mM sodium ascorbate (A4034, Sigma-Aldrich), 5 mM Trolox (238813, Sigma-Aldrich), and 0.02% sodium azide to inactivate the photochemical reaction. Cells were then washed three times with PBS-T.

Labeling validation

Labeled molecules were stained with NeutrAvidin-DyLight 488 (22832, Thermo Fisher Scientific) for 1h in 3% BSA/PBS-T. The antibody-$[Ru(bpy)_3]^{2+}$ conjugates were hybridized with a fluorescence marker (goat anti donkey TexasRed, PA1-28736, Thermo Fisher Scientific) for 1 h at RT. Cells were subsequently stained with nuclear marker (Hoechst 33258, 62249, Thermo Fisher Scientific) for 30 min at RT. A confocal microscope (LSM 880, Zeiss, Germany) was then used for 3D imaging.

Protein extraction and on-bead digestion

Labeled cells were harvested by scraping with buffer containing 10 mM Tris (pH 8.0), 1% Triton X-100, 1-fold protease inhibitor cocktail, 10 mM sodium ascorbate, 5 mM Trolox, and 1 mM sodium azide.  Harvested cells were sonicated using a Q125 sonicator (Qsonica, CT, USA) and then subject to evaporation of the scraping buffer by SpeedVac (Concentrator Plus, Eppendorf, Germany). Lysis buffer containing 4% sodium dodecyl sulfate (SDS), 1% Triton X-100, 10 mM

Tris (pH 8.0), and dithiothreitol (DTT) were added to the harvested cells. To retrieve the crosslinked amide groups resulted from PFA fixation, the lysed cells were further heated at 99°C for 45 min. Lysates were centrifuged and the supernatant was collected. Streptavidin magnetic beads (88817, Thermo Fisher Scientific) were washed with a dilution buffer (0.5% Triton X-100/PBS) for three times, and the protein lysates were diluted 10-fold to reduce the SDS concentration to less than 0.4%. The diluted lysates were added to the washed beads and incubated at 2-8°C for 16 h with rotation. The biotinylated-protein bonded beads were washed to remove non-specific binding. For on-bead digestion, the beads were further washed with 100-μL 50-mM TEABC (T7408, Sigma-Aldrich) for three times, and the biotin-protein bonded beads were then mixed with 0.2 μg of trypsin/ Lys-C (V5071, Promega, WL, USA) to a final volume of 20 μL at 37°C for 100 min for an initial digestion. The supernatant was collected and subject to overnight digestion. Finally, the digests were acidified by adding 2 μL of 10% formic acid and desalted by C18 Ziptip (ZTC18S096, Merck, MA, USA). Desalted peptides were dried by Speedvac and stored at -20°C prior to LC-MS/MS analysis.

LC-MS/MS analysis

Detection of immunoprecipitated product was done by data-dependent acquisition mass spectrometry. LC-MS/MS analysis was performed using an UltiMate 3000 RSLCnano system (Thermo Fisher Scientific) coupled to an Orbitrap Fusion Lumos mass spectrometer (Thermo Fisher Scientific). The desalted peptides were resuspended in 0.1% formic acid in water and loaded onto a PepMap™ 100 C18 HPLC column (2 µm, 100 angstrom, 75 µm × 25 cm; Thermo Fisher Scientific), and peptides were eluted over 160-min gradients for nuclei-illuminated samples, over 120-min gradients for nucleoli-, stress granule-illuminated samples. The full MS spectra ranging from m/z 375–1500 were acquired at a resolving power of 120,000 in Orbitrap, an AGC target value of $4 \times 10^5$, and a maximum injection time of 50 ms. Fragment ion spectra were recorded in the top-speed mode at a resolving power of 30,000 in Orbitrap using a data-dependent method. Monoisotopic precursor ions were selected by the quadrupole using an isolation window of 1.2, 0.7, 0.4 Th for the ion with 2+, 3+, 4–7 charge states, respectively. An AGC target of $5 \times 10^4$, maximum injection time of 54 ms, higher-energy collisional dissociation (HCD) fragmentation with 30% collision energy, and a maximum cycle time of 3 s were applied. Dynamic exclusion was set to 60 s with an exclusion window of 10 ppm. Precursor ions with the charge state of unassigned, 1+, or superior to 8+ were excluded from fragmentation selection.

Protein identification and label-free quantification

Raw data from the same batch of two-photon illumination were processed with Proteome Discoverer (Thermo Fisher Scientific) by Sequest HT algorithm against the UniProtKB/Swiss-Prot human protein database (version 2020.02, 20,365 entries) for feature extraction, peptide identification, and protein inference. Database search was performed with the following criteria and parameters: tryptic peptides with up to three missed cleavages; mass tolerances of 10 ppm for peptide ions, and 0.05 Da for fragment ions; static carbamidomethylation (+57.0215 Da) on Cys residues; dynamic deamidation (+0.9840 Da) on Asp and Gln residues, oxidation (15.9949 Da) on Met residues, acetylation on protein N-termini (+42.0106 Da), and desthiobiotin phenol modification (+331.1896) on Tyr residues. The minimal peptide length was set to 6 residues. The false discovery rate (FDR) of peptide and protein were both set to 1%. For label-free quantification, the time window for chromatographic peak alignment was set to 20 min. Peptide level data was then normalized to the total peptide intensity, and the quantification value for a given protein was derived from the sum of normalized intensities of the top three intense unique peptides belonging to that protein. Label-free quantification (LFQ) was performed to extract the enriched proteome from background signals. After streptavidin bead enrichment and mass spectrometry identification, protein identified from both the control group and the experimental group were analyzed by comparing their peak intensities. The distribution of true positive and false positive proteins was separated in the axis of $\log_2$(PL/Control) and the photolabeled proteome was reported by applying a cutoff of $\log_2$(PL/Control).

Streptavidin dot-blot analysis

Dot-blot analysis was performed on a PVDF membrane. PVDF membrane was activated with 100% MEOH and soaked in PBS-T for 2 min. 8 µl of each sample were spotted on the membrane. After drying, the membrane (blot) was rinsed with PBS-T and blocked with 5% BSA for 30 min at RT, followed by a wash with PBS-T. The blot was then incubated with streptavidin-horseradish peroxidase (HRP, 1:1000) in 5% BSA at 4°C overnight. The blot was subsequently washed with PBS-T prior to the development with ECL substrate (Bio-Rad, CA, USA) using iBright FL1500 image system (Cytiva). For re-probing, the same blot was stripped three times with NaOH for 5 min at 60°C, and then blocked with 2.5% BSA, and re-probed with mouse anti-tubulin at a 1:500 dilution (GTX112141, GeneTex, CA, USA) in 2.5% BSA at 4°C overnight, and washed with PBS-T. The blot was incubated with anti-mouse-HRP (1:500) in 2.5% BSA for 2 h at RT. The blot was washed with PBS-T prior to the development with ECL substrate using iBright FL1500 imaging system.

Functional annotation

The GO term association was obtained from Uniprot and the protein complex data was downloaded from CORUM. The Fisher's exact test with the Benjamini-Hochberg multiple testing correction was used to identify over-represented GO terms and protein complexes. The proteins annotated by GO and curated by CORUM were used as the background set for GO and protein complex enrichment analyses. STRING analysis of stress granules protein network was performed using 124 over-represented (enriched) proteins for associations regarding functional and physical interactions as shown in a visualized network graph.

Colocalization validation

For the stress granule study, the arsenite induced or control U-2OS cells were fixed with 2.4% PFA or MEOH as described above. The cells were then incubated overnight at 4°C with primary rabbit antibodies against PDLIM7 (PA5-53846, Thermo Fisher Scientific), EIF3CL (ab237757, Abcam), YWHAE (ab92311, Abcam), RPSA (MA5-32499, Thermo Fisher Scientific), MTA2 (ab171073, Abcam), UGDH (PA5-88347, Thermo Fisher Scientific), DDX17(DF12935, Biosciences), ANLN (PA5-114851, Thermo Fisher Scientific), PSMD3 (PA1-974-A488, Thermo Fisher Scientific), PSMA6 (PA5-76058, Thermo Fisher Scientific), MCM2 (PA5-78054, Thermo Fisher Scientific), and PPIA (ab3563, Abcam). Cells were washed three times with PBS-T, and then incubated with stress granule markers mouse anti-G3BP1 (sc-365338, Santa Cruz) for 2 h at RT. Cells were then stained with secondary AlexaFluor 488-goat anti-rabbit IgG (A21245, Thermo Fisher Scientific) and AlexaFluor 568-goat anti-mouse IgG (A10037, Thermo Fisher Scientific) followed by Hoechst staining (62249, Thermo Fisher Scientific). For the amyloid-β study, Aβ1-42 plaque expressed SH-SY5Y cells were fixed with 2.4 % PFA, and then incubated overnight at 4˚C with primary rabbit antibodies against Lon protease (H00009361-D01P, Abnova, Taiwan), and DDX3X (PA5-17165, Thermo Fisher Scientific) and primary mouse antibody against Aβ (Aβ1-16, 803014, BioLegend, CA). Cells were washed three times with PBS-T, and then incubated with secondary AlexaFluor 647-donkey anti-rabbit IgG (A32795, Thermo Fisher Scientific) and AlexaFluor 555-goat anti-mouse IgG (A32727, Thermo Fisher Scientific) followed by DAPI staining (D9542, Sigma-Aldrich). All images were acquired by confocal microscopy (LSM 880, Zeiss).
